# Benchmarking of tools for axon length measurement in individually-labeled projection neurons

**DOI:** 10.1101/2021.05.11.443544

**Authors:** Mario Rubio-Teves, Sergio Diez-Hermano, César Porrero, Abel Sánchez-Jiménez, Lucía Prensa, Francisco Clasca, María García-Amado, José Antonio Villacorta-Atienza

## Abstract

Projection neurons are the commonest neuronal type in the mammalian forebrain and their individual characterization is a crucial step to understand how neural circuitry operates. These cells have an axon whose arborizations extend over long distances, branching in complex patterns and/or in multiple brain regions. Axon length is a principal estimate of the functional impact of the neuron, as it directly correlates with the number of synapses formed by the axon in its target regions; however, its measurement by direct 3D axonal tracing is a slow and labor-intensive method. On the contrary, axon length estimations have been recently proposed as an effective and accessible alternative, allowing a fast approach to the functional significance of the single neuron. Here, we analyze the accuracy and efficiency of the most used length estimation tools - design-based stereology by virtual planes or spheres, and mathematical correction of the 2D projected-axon length - in contrast with direct measurement, to quantify individual axon length. To this end, we computationally simulated each tool, applied them over a dataset of 951 3D-reconstructed axons (from NeuroMorpho.org), and compared the generated length values with their 3D reconstruction counterparts. Additionally, the computational results were compared with estimated and direct measurements of individual axon lengths performed on actual brain tissue sections, to analyze the practical difficulties and biases arising in real cases. The evaluated reliability of each axon length estimation method is then balanced with the required human effort, experience and know-how, and economic affordability. This work, therefore, aims to provide a constructive benchmark to help guide the selection of the most efficient method for measuring specific axonal morphologies according to the particular circumstances of the conducted research.

**AUTHOR SUMMARY:** Characterization of single neurons is a crucial step to understand how neural circuitry operates. Visualization of individual neurons is feasible thanks to labelling techniques that allows precise measurements at cellular resolution. This milestone gave access to powerful estimators of the functional impact of a neuron, such as axon length. Although techniques relying on direct 3D reconstruction of individual axons are the gold standard, handiness and accessibility are still an issue. Indirect estimations of axon length have been proposed as agile and effective alternatives, each offering different solutions to the accuracy-cost tradeoff. In this work we report a computational benchmarking between three experimental tools used for axon length estimation on brain tissue sections. Performance of each tool was simulated and tested for 951 3D-reconstructed axons, by comparing estimated axon lengths against direct measurements. Assessment of suitability to different research and funding circumstances is also provided, taking into consideration factors such as training expertise, economic cost and required equipment, alongside methodological results. These findings could be an important reference for research on neuronal wiring, as well as for broader studies involving neuroanatomical and neural circuit modelling.

## INTRODUCTION

The highly integrated functioning of the brain relies on axons that directly connect distant regions. The introduction in the past decade of new viral vectors able to drive the in vivo expression of high levels of marker proteins has allowed, for the first time, the consistent and complete visualization of such long-range projection axons with single-cell resolution [1–6]. Studies in small rodent brains applying these methods have revealed that despite being submicron-thick, an individual neuron axon can extend over long distances and branch in complex and specific patterns [6–9].

Unlike dendrites, which integrate calcium signals in mainly linear fashion [10, 11], the axon is a transmission compartment for all-or-none fast sodium axon potentials. The functional impact of signals travelling down an axon thus depends critically on the wiring of the axon, as it constrains the number and distribution of its synapses. Axon morphologies may in this way lead to different network configurations, so precise axon characterization is a must for modelling and describing a plethora of brain capabilities [12–14]. Interestingly, the number and distribution of synapses can be reliably derived from the axon length within its target structures [8, 9, 15]. Accurate measurement of axon length, therefore, is key for the functional modelling of brain-wide circuits as it estimates the functional impact of the single neuron [16, 17].

However, mapping and measuring long-range projection axon trees remains challenging, as it requires working across a wide range of spatial scales, from submicron to brain-wide [18]. A significant recent advance has been the development of automated platforms that combine serial sectioning with high-resolution confocal image acquisition to produce massive whole-brain volumetric image datasets. On these datasets, fluorescently tagged axons can be traced and measured using tools for 3D navigation and annotation. Deformation of these datasets is known and measurable, and the axon length measurements are thus reliable as far as the reconstructions are complete. Nevertheless, devices for producing high-resolution 3D datasets are prohibitively expensive and the resulting multi-terabyte datasets require high-end computing infrastructure. For the foreseeable future, these high-end platforms will remain limited to industrial-scale research facilities. At present they remain focused in producing open-access datasets of adult “standard” model species brains intended as reference for other brain circuit studies [6, 19–21].

Hundreds of laboratories and imaging facilities around the world currently rely on small, camera-lucida (2D) or computerized (3D) systems to reconstruct and measure neuron morphologies from brain tissue sections [22–24]. These systems remain the feasible way for reconstructing neuronal morphologies on brains manipulated under specific experimental conditions, or when in situ visualization of specific tissue markers is required, in developing animals or in species other than mice. However, producing accurate axon length data from serial histological sections is not trivial, as projection axons can extend across the brain and thus be spread over a large number of tissue sections. Thus, it is important to evaluate the accuracy and efficiency of the procedures currently in use, with the aim of comparing data available in the literature from different methods.

Direct measurements of axon length can be obtained through three-dimensional (3D) reconstruction-tracing of arborizations across serial sections [25]. This is usually done manually and is slow and work-intensive, as it must be done at high magnification and deals with uneven distortions and misalignments due to histological tissue processing. A widely used computerized tool is Neurolucida (MBF Bioscience, Williston, VT, USA), which allows for manual tracing among other capabilities, but a wide array of software tools from different sources is also available online [23]. Regardless of how sophisticated the software used is, the basis for length estimation is still the same: transforming image data into a series of vectors with a tree-like structure defined by a set of parameters that includes its euclidean coordinates, from which an approximation to the real, total path length is derived.

A faster, indirect alternative is to produce a 2D reconstruction by projecting onto a plane using a camera lucida or a slide-scanner, and then multiplying it by a correction factor [2, 7]. Although this method has been recently applied to a wide range of thalamocortical neurons, the accuracy of the estimation method and the suitability of the correction factor has not been tested before.

Other indirect approaches to measure axon length are based on stereological techniques. Design-based stereology takes advantage of unbiased sampling schemes with different kinds of probes (planes or spheres) that are applied to the tissue to produce a number of interactions with the object of interest (the axon) that, because of the randomness of the sampling, can be used to mathematically infer its length with a known error [26–28]. These stereological methods have been traditionally applied to estimate the axonal length of neuronal populations ([29, 30] for virtual planes; [31–33] for virtual spheres) and, more recently, of individual neurons [34].

In the present study, we sought to compare these estimation tools when measuring individual highly complex axonal arbors. This way, we propose a twofold objective. On the one hand, we assess the accuracy and efficiency of these methods to provide a rigorous rationale of their pros and cons. On the other hand, we evaluate their performance when estimating individual axon length and so as a qualitative approach to the functional importance of a single neuron. Taking advantage of the large number of neuron morphologies available at NeuroMorpho.org [35], we computationally implemented length estimation protocols from model-based estimates (mathematical correction of 2D length) and design-based stereology (sampling with isotropic virtual planes and spheres) to compare among themselves and with direct reconstruction-derived measurements. Our results provide: 1) a classification of the different estimation methods according to their accuracy and performing effort, 2) a guide for selecting the most suitable parameters for each length estimation method in line with the researcher’s necessities (type of axon characterization demanded by the problem, available funding and personnel, assumable error, time to be invested, etc.) and 3) a summary of advantages and disadvantages of estimates vs. direct methods.

## MATERIAL AND METHODS

### Neuron sample

A sample of 951 neuron morphology files was downloaded from the NeuroMorpho database (NeuroMorpho.Org, RRID:SCR_002145; [35].

We searched the database for full-morphology files produced on the mouse model (*Mus musculus*). The neurons contained in those files had been uploaded to NeuroMorpho.org by a number of different laboratories, and were produced either through direct reconstruction on a series of sections from a sparsely-labeled brain (containing one or a few neurons) that spanned the total length of the arbor or, in the case of those belonging to the MouseLight project, through semiautomatic tracing of single cells from a densely labeled (containing hundreds of neurons) brain digitally scanned by means of single two-photon tomography (STPT) [4]. Although different in practice, these two technical approaches are conceptually the same and were considered equivalent in the coming analysis. That is, the axonal length of the 3D reconstructions was considered as the “true value” and used to measure the accuracy of the different estimations.

Neurons were then qualitatively classified into four distinct classes based on easily recognizable features of their morphology and previous classifications (**Fig. 1**; [36]): type 1 (“specific”) neurons were characterized by a single focal arborization (N = 90); type 2 (“multispecific”) neurons displayed focal clusters in multiple spatial locations (N = 116); type 3 (“non-specific”) neurons featured axons that never developed dense, focal, highly-ramified clusters, but instead had a much more sparse or scattered appearance (N = 269); lastly, type 4 (“local”) neurons resembled interneurons of different shapes and sizes and were, because of their smaller size and ease of reconstruction, the more numerous in the dataset (N = 476). Sometimes neurons had characteristics that could cause them to be assigned to more than one category; in those cases, the more predominant feature decided what type it belonged to. This classification pays no attention to the anatomical position or developmental origin of each cell, and was performed previous to the analysis in order to see if gross morphological appearance could be used to infer the applicability of a specific method to a particular neuron.

**Figure 1.**
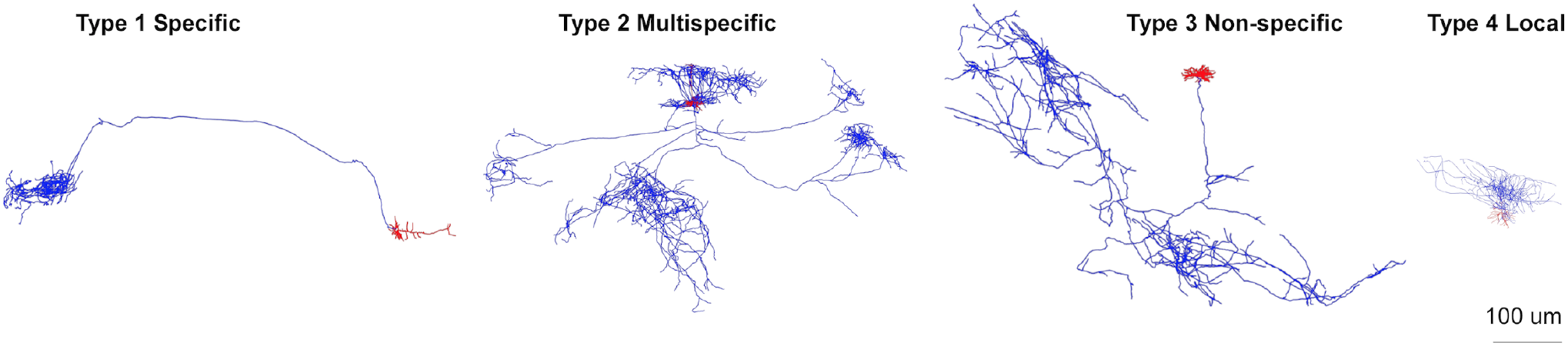
Neuron classification. Representative examples of neurons from the NeuroMorpho dataset showing the four axonal types considered in the study (Drawings generated from morphologies AA0641, AA0002, AA0051, AA0771 from Neuromorpho.org).

### Computational implementation of axon length estimation methods

The full-morphology files from the database were analyzed by means of custom-made Matlab scripts (Matlab 2017b; The Mathworks, Natick, MA, USA), where only the information for the axon was treated. Projection-based method was implemented by eliminating the Z coordinates from the axon points to project them onto the XY plane. The length of the resulting projected axon multiplied by a proper coefficient (see Results) will then be used as estimation of the axon tridimensional length. Note that the XY plane was not *a priori* established but defined by its own morphology file and each source’s method of acquisition. Sphere and plane-based estimation methods were implemented following similar processes. The analyzed axon was divided into sections of 50 μm thick parallel to the XY plane (plane defined by X and Y coordinates in the morphology file). In both cases the sampling box dimensions were 50×50×50 μm^3^ and all sections were analyzed. The specific probes (spheres and planes) were introduced into the sampling box according to the requirements of each particular method (random orientation in each sampling box for plane-based method). The relevant parameters, as sphere diameter, plane distance, and step length between sampling boxes, were adjusted accordingly for the computational simulations (**Fig. 2**).

**Figure 2.**
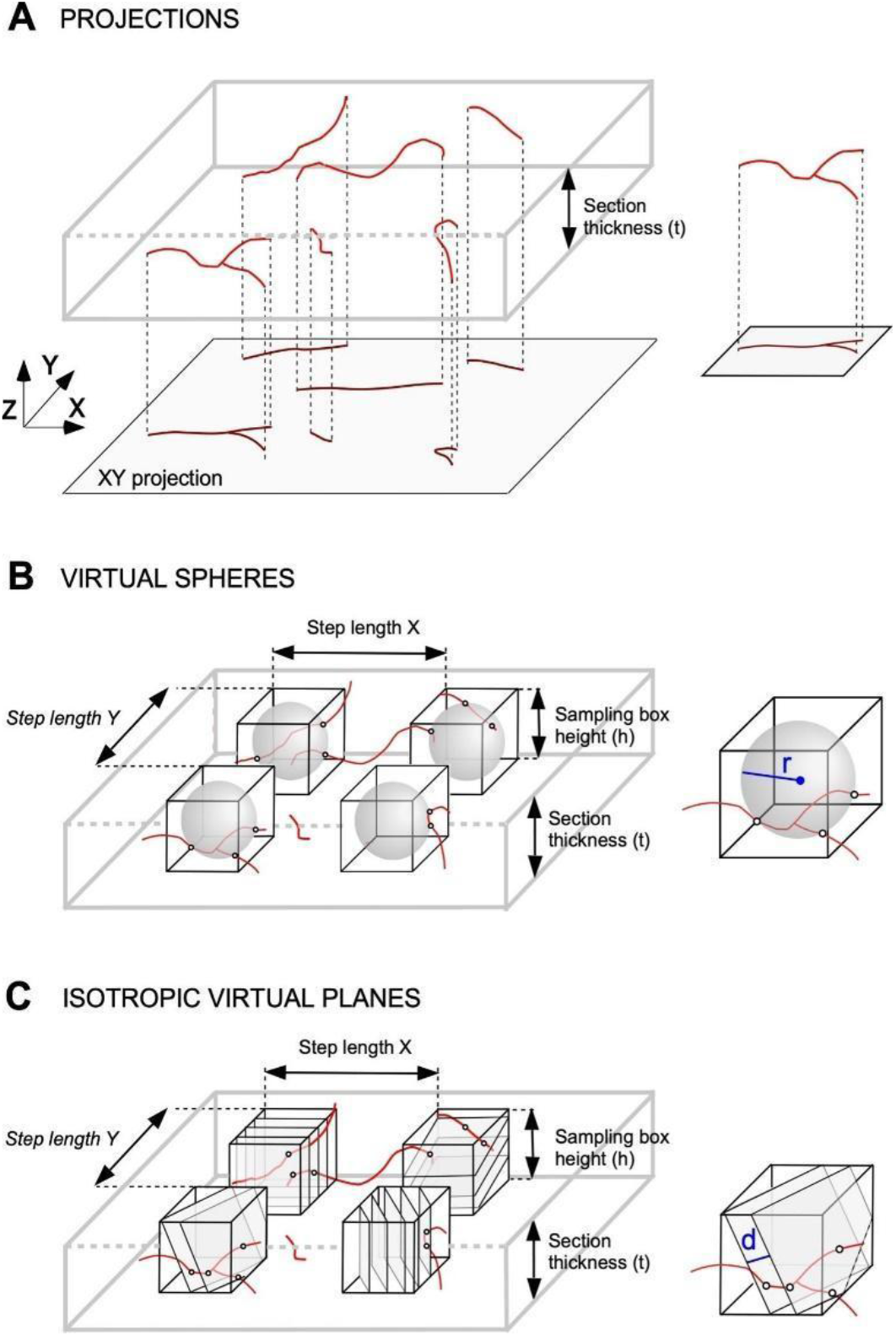
Methods for length estimation. **A**. Projections method. One tissue section containing several axon fibers (above). The method consists of measuring the axon length projected in the X-Y plane (represented below) and multiplying it by a factor to obtain the 3D actual length. **B**. Stereology with virtual spheres. This method is based on the estimation of axonal length from the intersections occurring between the axon segments and the surface of virtual spheres introduced in sampling boxes inside the tissue section thickness. Four sampling boxes inside a section are represented in black, with the virtual spheres inside (in grey). **C**. Stereology with virtual planes estimates axonal length by using the intersections between the axon segments (in red) and a set of parallel and isotropically oriented planes contained within the sampling boxes included in the tissue sections. Four sampling boxes (in black) with parallel virtual planes inside (in grey) are represented; r and d stand respectively for sphere radius and distance between planes.

### Statistical analysis

#### Projection-based length estimation

Correlation between axon’s real length and projection’s length was assessed by means of linear regression (ordinary least squares, OLS), with axon type considered as a factor. Parameter estimators (alphas) were obtained through error-resampling bootstrapping (2000 repetitions). 95% confidence intervals (CI95%) were calculated using the 0.025 (lower limit) and 0.975 (upper limit) quantiles of the bootstrapped distribution of alpha values. Prediction errors were estimated through 5-fold cross validation (CV), independently for every axon type and plane combination. Error’s density distributions were obtained through bootstrapping (100 repetitions), and probabilities were calculated as the area under the curve for a given error interval (i.e. + 5% and +10%).

#### 3d plane-based stereology

Model errors were calculated as the absolute difference between axon’s real length and length estimated by all combinations of sampling step and distance between planes. Correlation between model error, step and distance was assessed by means of linear regression (OLS), independently for every axon type. Prediction errors were calculated as smoothed interpolations for 200 possible values of step and distance. Error’s density distributions were obtained as smoothed histograms with automatic bandwidth selection, and probabilities were calculated as the area under the curve for every step and distance combination and for a given error interval (i.e. + 5% and +10%). Additionally, all models defined by step and distance were compared in terms of normalized root mean square error (RMSE) by means of 5-fold CV.

#### 3d sphere-based stereology

Analysis followed the same procedures as the plane-based stereology, considering sampling step and probe diameters in this case.

Mean absolute error, mean intersections, and error probability distributions were obtained through linear interpolation from these estimations performed for diameter values in [10, 15, 20,…, 50] (virtual spheres), distance values in [3, 6, 9,…, 30] (virtual planes) and step values in [70, 80,…, 150] (both probe types). The adjusted R-squared coefficients range from 0.88 to 0.98.

### Practical implementation

We performed practical examples with real tissue sections comparing the three methods analyzed in the computational approach: direct axon measurements from Neurolucida reconstructions, projections-based length estimation and design-based stereology (virtual planes) (**Fig. 2**).

We measured the axonal length of three mouse neurons which fitted in categories 1, 2, and 3 of our study by using the mentioned three length estimation methods (**Fig. 3**). We did not include type 4 neurons in the experimental approach based on the results obtained with the computational approach (see Results section). Experimental procedures including mice surgery, virus injection for labelling neurons and immunohistochemistry as previously described [5]. All procedures involving live animals were conducted under protocols approved by the University ethics committee and the competent Spanish Government agency (PROEX175/16), in accordance with the European Community Council Directive 2010/63/UE. The soma of these neurons was located in the thalamus, and their axon extended to different targets in the cerebral cortex. The complete axon inside the cortex was reconstructed using the Neurolucida software (Neurolucida 2020; MBF Bioscience, Williston, VT, USA) and the length obtained with this direct measurement was compared with the resulting axonal length from both the use of virtual planes (stereology) or the projections method. We used 50 microns-thick sections containing the complete axon of each neuron.

**Figure 3.**
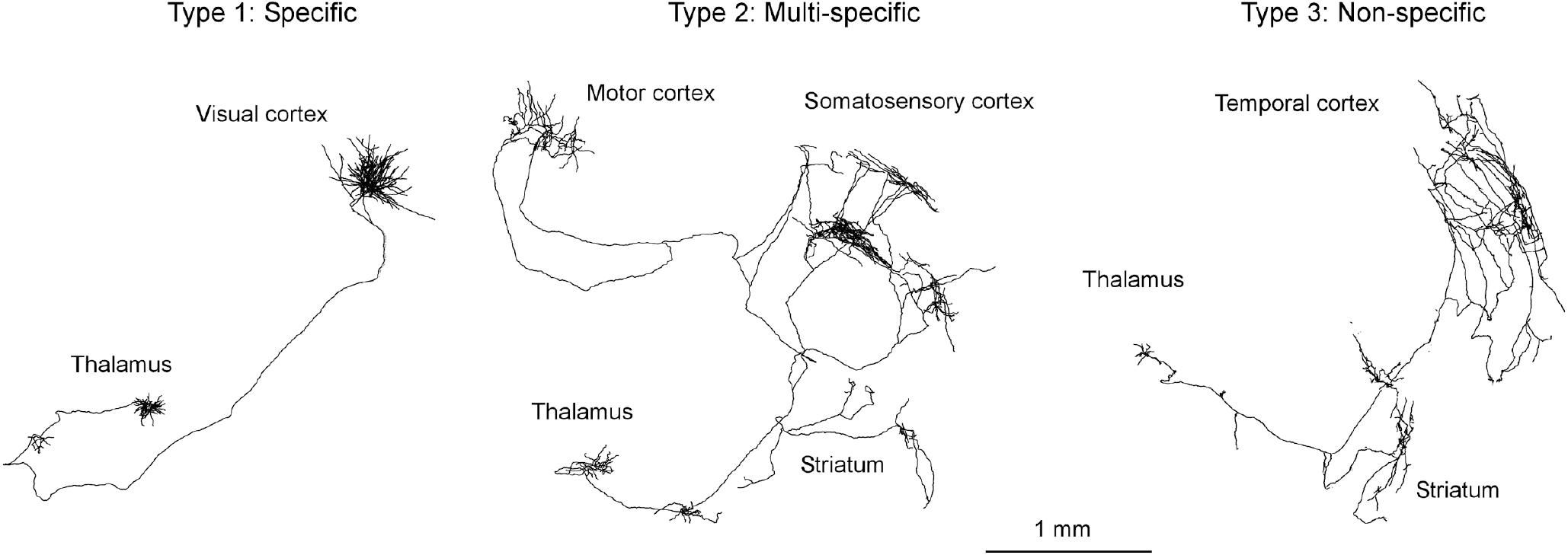
Complete reconstruction of the three mouse neurons used for the practical implementation of the length estimations methods in the laboratory. For all the cases, the soma was located in the thalamus and the axon reached different regions of the cerebral cortex.

The projection-based method consisted in drawing the complete axon of each neuron, contained in the corresponding set of sections, by using the 20X lens of a microscope connected to a camera lucida (Nikon Eclipse E400; Nikon, Tokyo, Japan). Camera lucida drawings containing the projected axon were scanned and redrawn for digitization on Canvas X GIS (Canvas GFX, Boston, MA, USA); this software was also used to extract the length of all axon segments. Projected length was multiplied by the proper alpha value of each axon type (see **Fig. 4B** and **“Projected based estimation” section**) to estimate the real axonal length.

**Figure 4.**
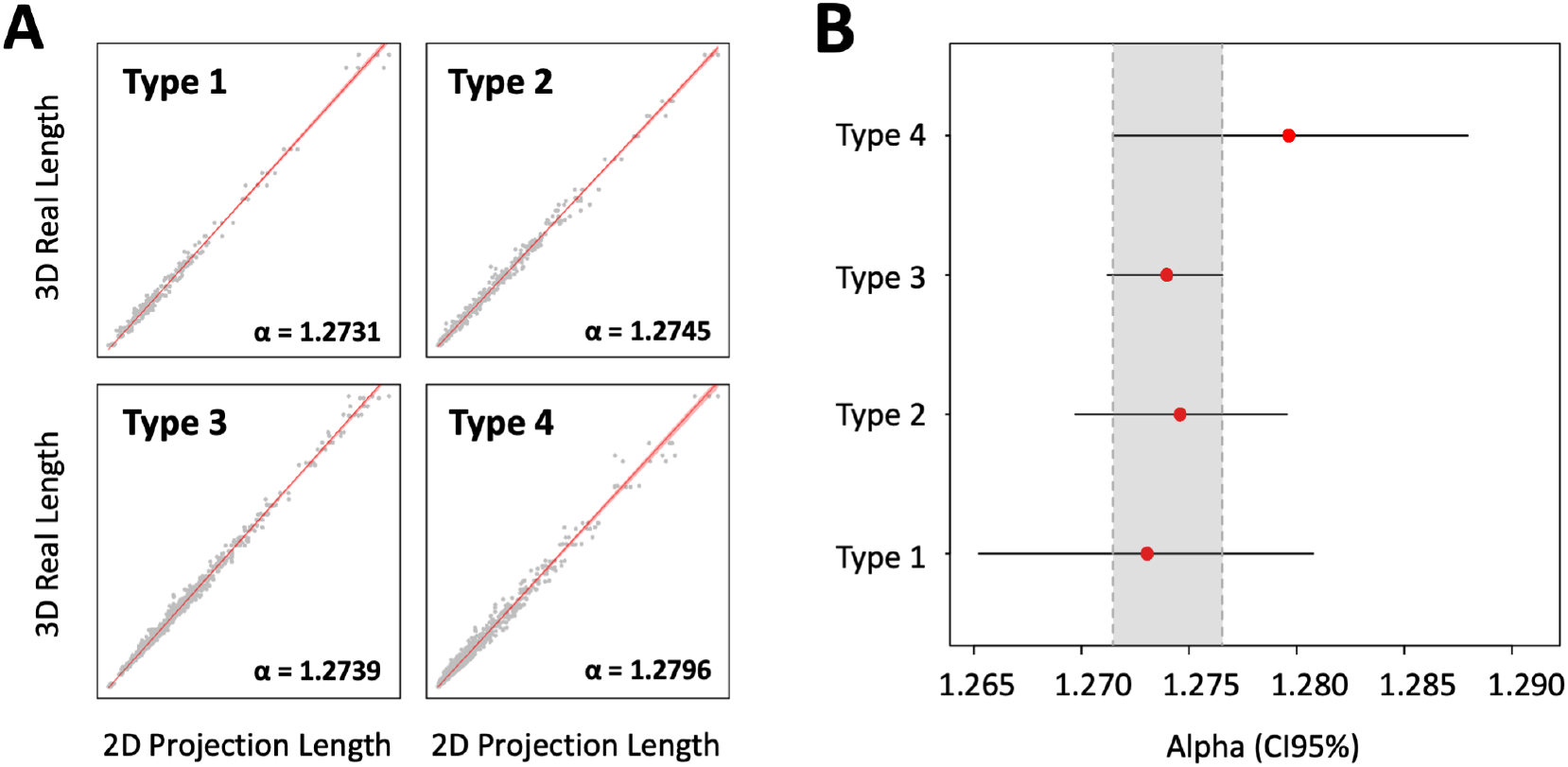
Axon length correlations for projection-based stereology. **A**. Correlation between 3D real length and 2D projection length, by axonal type. Red lines represent the mean estimated 3D real length (ordinary least squares), and light red shaded areas correspond to 95% confidence intervals (CI95%). Gray dots depict full samples comprising all planes measurements (XY, XZ and YZ, note the organization of the data in triplets). Global adjusted R-squared = 0.99. **B**. Estimated slopes (red dots) and CI95% (horizontal black bars) values for correlations in A. Slopes can be interpreted as coefficients (alpha) that multiply 2D projection values to obtain predictions of 3D real lengths. Note that all axon types share a common set of potential alphas at a 95% confidence level (gray shaded area).

The stereological approach consisted on projecting virtual planes over the sections observed in a microscope connected to a computer running stereology software (newCAST stereology software package for VIS; Visiopharm, Denmark); sampling parameters used for the 3 neuron types were: 75 μm step length in X and Y axes; sampling box size: 50 (X axis), 50 (Y axis) and 10 (Z axis) μm and 5 μm of plane separation; a fractionator sampling scheme was used to estimate the axonal length [37]. We calculated the coefficient of error (CE) due to the sampling method for the estimates by using the equations detailed in [38].

## RESULTS

### Projection-based estimation

The projection-based method of length estimation relies on the assumption of isotropy (that fibers distribute their length equally across all possible directions; [22]). Under this condition the 3D length of isotropic fibers is equal to the product of the two-dimensional (2D) length of its projection onto a plane by a constant factor of π/2 (**Fig. 2A**). However, neurons do not distribute their fibers randomly, so the accuracy of this estimation will ultimately depend on the degree of anisotropy of the cell it is being applied to. Another way of seeing this is that each neuron will have its own correction factor, depending on its degree of anisotropy.

In order to analyze the reliability of projection-based axon length estimation in real neurons, we projected the morphologies of the dataset from NeuroMorpho.org onto the XY, XZ and YZ planes. We found that the relationship between the 2D projection length and the reference length of the 3D model is indeed linear for all four types of neurons (**Fig. 4A**). This confirms there is a coefficient describing the relationship between the 3D length of the real structure and the length of its projection onto a plane. For the dataset used in this study, we calculated the alpha values for the different types of axons in our classification, whose 95% confidence intervals (95% CI) overlap between 1.272 and 1.277 (**Fig. 4B**). The particular alpha values for each axon type are detailed in **Fig. 4A**.

Next, we quantified the probability of estimation error, defined as the difference between the 3D axon length and the length estimated from the model, when the different alpha factors are applied to the corresponding axon type in the projection-based approach. In order to do that, all axons were projected onto XY, XZ and YZ planes and their 3D length were estimated by multiplying the projection length by alpha. The area under curves in **Fig. 5A** describes the probability of finding a specific estimation error; as an example, dark and lightly shaded areas correspond to the probability of getting an estimation error of 5 and 10%, respectively. Noticeably, these cover much of the area under the curve, which means that it is highly probable to get estimation errors lower than 10% when using this approach. Specifically (**Fig. 5B**), types 1, 2, 3 and 4 neurons had a 65, 78, 71 and 38% probability of having a 5% absolute estimation error (estimation error between −5% and 5%), whereas the probability of it being 10% was 88, 92, 88 and 62%, respectively. It should be remarked that these estimation errors were similar across orthogonal projection planes, showing the robustness of the method when applied in practice since the projection plane is not a relevant parameter.

**Figure 5.**
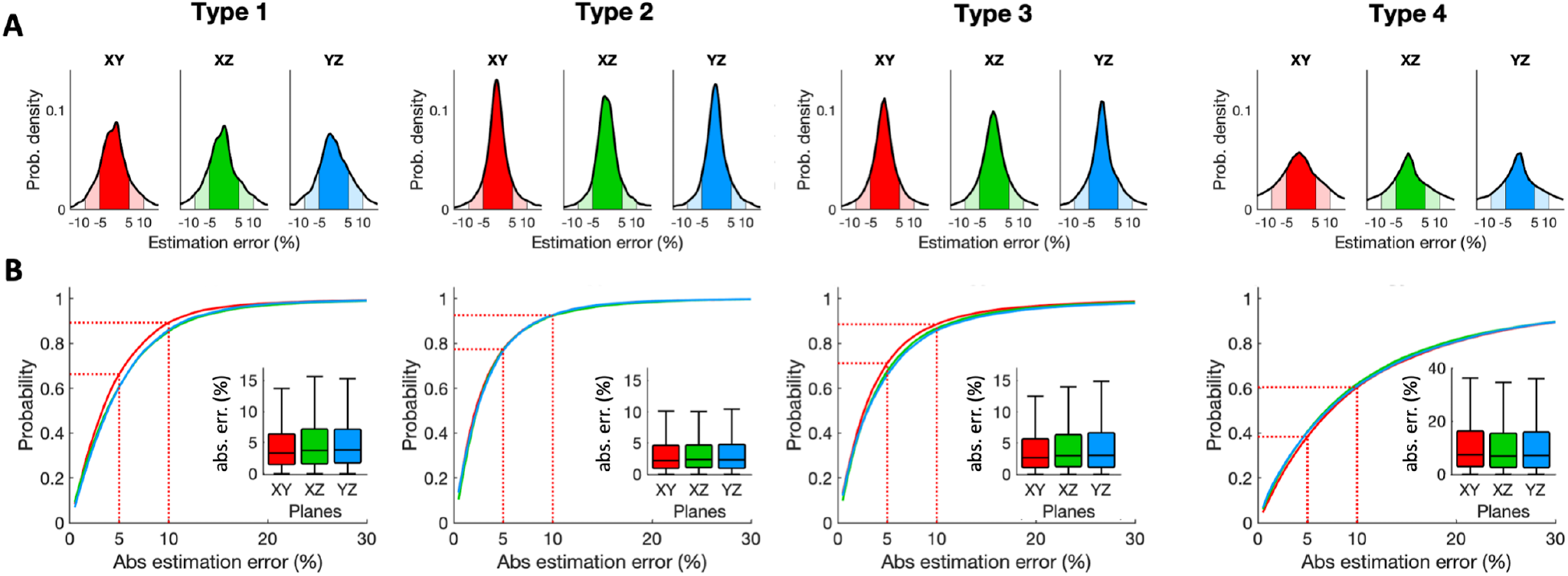
Estimation error probability projection-based stereology. **A**. Estimation error is distributed according to the shown probability densities (axon types in columns; red, green, and blue for XY, XZ, and YZ planes respectively). The probability of getting a certain error during axon length estimation will be given by the area below the curves (dark and light colors for + 5% and + 10% estimation error respectively). **B**. Absolute estimation error vs. probability (given by the area below the curves in A). Dashed lines point the probabilities of estimating axon length with absolute errors of 5% and 10%, (colored areas in A). Insets. Absolute error distributions for planes XY, XZ, and YZ.

### Stereological sampling with virtual spheres to estimate 3D length

In order to attain an unbiased stereological estimate, every object (whose length will be estimated) in the specimen has to have the same probability of being sampled. In the case of length estimates, 2D probes (surfaces) are used to scan the sample volume. The object length is thus estimated by quantifying the intersections between the object and the probe, whose position and orientation must be random to ensure an unbiased estimation. The stereological sampling scheme [27] draws on the inherent advantages of virtual spheres to use them as probes in systematic random sampling schemes aimed at achieving length estimates (**Fig. 2B**). The intersections between the isotropic surface of the sphere with the axons are counted, and the resulting number is used to estimate the total length under the conditions of randomness and isotropy above stated. The relevant parameters used in this estimation are the step distance (distance between the spheres in X and Y axes, which determines the number of spheres), and the sphere diameter (which determines the sampling surface dimensions).

To evaluate the efficiency of this method to estimate axon length, we performed sampling with virtual spheres on the cells in the dataset using different combinations of step distance and sphere diameter (**Fig. 6**). The results show that the mean absolute error in the length estimation when using this approach was surprisingly high, reaching values of 60-70% across all types (**Fig. 6A**). In neuron types 1, 2 and 3 the mean absolute error mainly depends on the probe diameter, being nearly independent of the step size. This is a consequence of using small spheres compared with the axon length. On the contrary, in type 4 (local) neurons, whose axons are significantly smaller than the other types (**Fig. 1**), the mean absolute error in the axon length estimation showed a dependence on both the sphere diameter and the step size.

**Figure 6.**
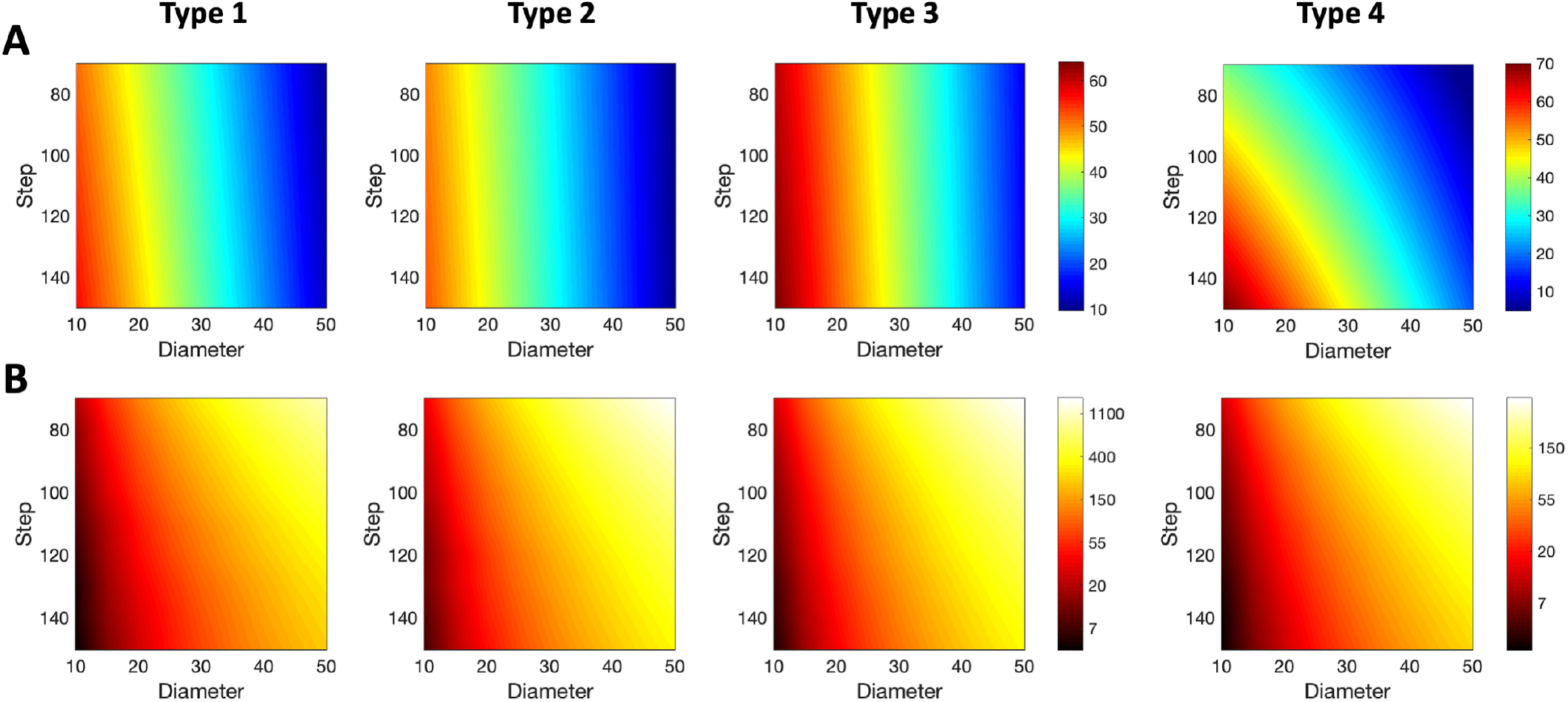
Mean absolute error for axon length estimation by spheres-based stereology. **A**. The mean absolute error (in % of the real axon length indicated by colors – see color bar) is shown for different values of the probe diameters and the step between sampling boxes, and for axon types 1 to 4 (columns). Note that the color bar is the same for Types 1, 2, and 3. **B**. Mean effort, quantized by the mean number of intersections, required to estimate the axon length with different values of the probe diameter and the step for axon types 1 to 4. Colors denote the number of intersections (in logarithmic scale; color bar is the same for types 1, 2, and 3). Mean intersection values were obtained through linear interpolation from estimation for diameter values in [10, 15, 20,…, 50] μm and step values in [70, 80,…, 150] μm.

In stereology the estimation error must be balanced with the sampling effort, which results from counting large numbers of intersections between the surface probe and the linear axon. Thus, we quantify the sampling effort directly as the number of intersections. **Fig. 6B** illustrates the mean effort of estimating axon length when using spherical probes, for the different types analyzed. It shows that for types 1, 2 and 3, achieving the lowest possible mean absolute error required counting between 200 and more than 1000 intersections, depending on the step size employed during the sampling (the higher the step, the lower the effort). This holds too for type 4 neurons, but the efforts required in general were significatively lower (50-250).

Therefore, **Fig. 6** is intended to estimate the parameter configuration according to the accuracy-effort trade-off. For instance, in case we would like to estimate the axon length of a type 1 single neuron by means of spheres-based stereology by only accepting a mean error lower than 10% and a mean effort lower than 300 intersections, matching **Figs. 6A** and **6B** would suggest the probe diameter equal to 50 and the step higher than 140.

### Stereological sampling with virtual planes to estimate 3D length

Other than spheres, it’s also possible to stereologically estimate the length of objects using isotropic (i.e., with random orientation) virtual planes as probes [26]. In this case, the sampling is performed in nearly the same way: sampling boxes containing the isotropic planes are spaced at a given XY step to randomly and systematically sample the tissue. The only difference is the introduction of the distance between planes inside the box instead of the diameter of the spheres as the second parameter critical for the stereological estimate (**Fig. 2C**).

The results of estimating axon length through sampling with isotropic virtual planes are depicted in **Fig. 7**. Depending on the combination of XY step and distances used, the mean absolute error for the estimation could be lower than 5% for types 1-3 and 10% for type 4. The best results were obtained when trying to maximize the sampling surface, that is, when combining small XY steps (80 μm) with small inter-plane distances (5 μm; **Fig. 7A**), at the cost of increasing the number of intersections.

**Figure 7.**
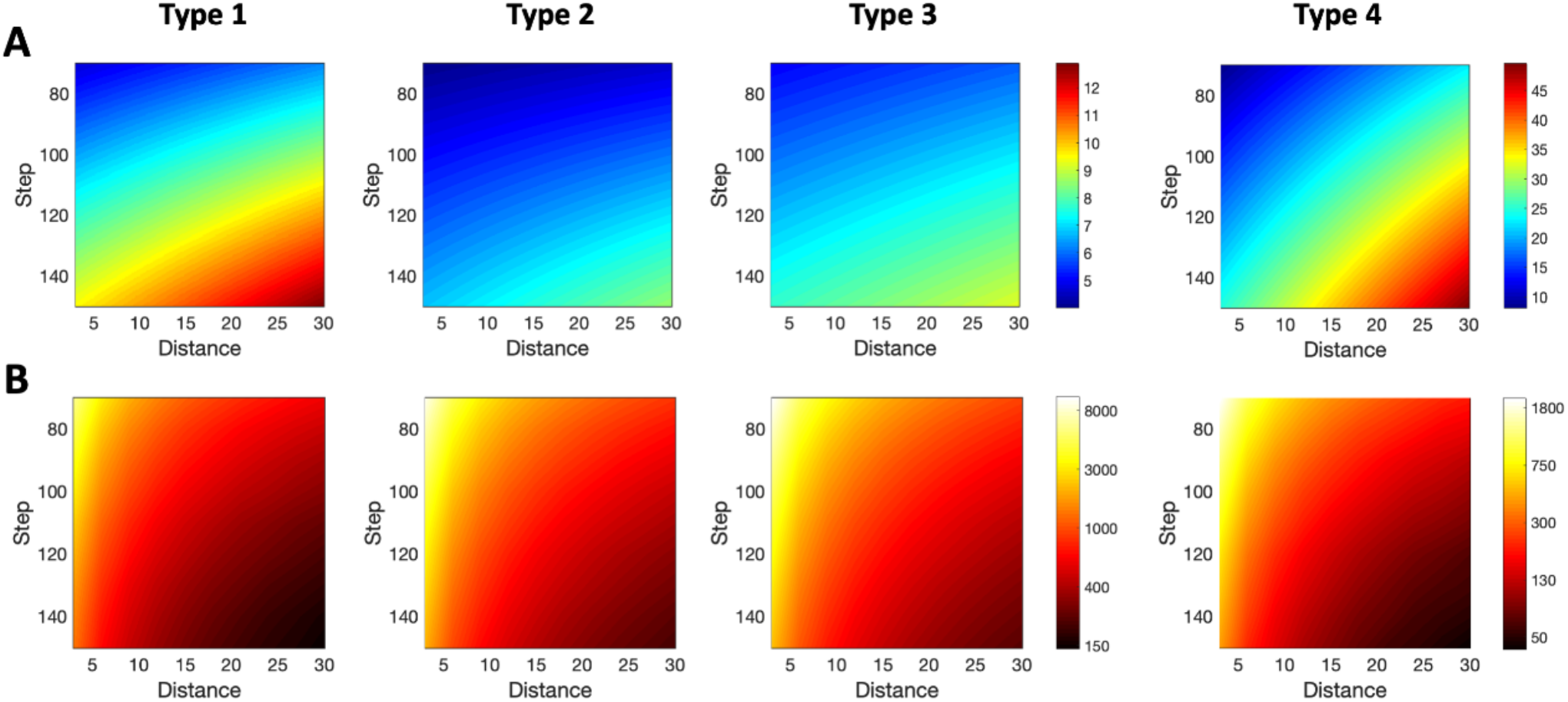
Mean absolute error for axon length estimation by planes-based stereology. **A**. The mean absolute error (in % of the real axon length indicated by colors – see color bar) is shown for different values of the distance between planes and the step between sampling boxes, and for axon types 1 to 4 (columns). Note that the color bar is the same for Types 1, 2, and 3. **B**. Mean effort, quantized by the mean number of intersections, required to estimate the axon length with different values of the distance between planes and the step for axon types 1 to 4. Colors denote the number of intersections (in logarithmic scale; color bar is the same for types 1, 2, and 3). Mean intersection values were obtained through linear interpolation from estimation for distance values in [3, 6, 9,…, 30] μm and step values in [70, 80,…, 150] μm.

The number of intersections was again used to quantify the effort demanded by the stereological procedure (**Fig. 7B**), since performing estimations with low mean absolute errors could require counting abnormally high numbers of intersections. However, specific combinations of parameters could also greatly reduce the mean effort while preserving an acceptable estimation error. For example, when estimating length for type-2 axons, small steps combined with high inter-plane distances produced similar errors at a significatively lower effort cost than small steps combined with small inter-plane distances. This way, by combining **Figs. 7A** and **7B**, a estimation of parameter configuration for plane-based stereology could be obtained according to the required mean error and effort.

Results in **Fig. 7** show the expected estimation error under different parameter combinations. Nonetheless, to make a right choice of XY step and plane distance, the researcher requires to know the probability of making a certain error under such parameters. This is detailed in **Fig. 8**, which shows the probability of attaining absolute estimation errors lower or equal to 5% (first row in **Fig. 8A**) and 10% (second row in **Fig. 8A**). To easily understand these distribution plots, **Fig. 8B** shows a particular example, denoted for each axon type by white dots in **Fig. 8A**. This example corresponds to a distance = 18 *μ*m and step = 110 *μ*m, and the two areas under each curve quantify the absolute error probability under or equal to 5% and 10% (color areas coincide with the color codes for the corresponding white dots). For instance, for type 1, error probability is equal to 0.39/0.68 (upper/lower panels in A; blue/green areas in first panel in B). That means if the researcher would estimate 100 samples with distance = 18 *μ* and step = 110 *μ*, in 35/68 samples the absolute error between real and estimated axon length will be below 5%/10%.

**Figure 8.**
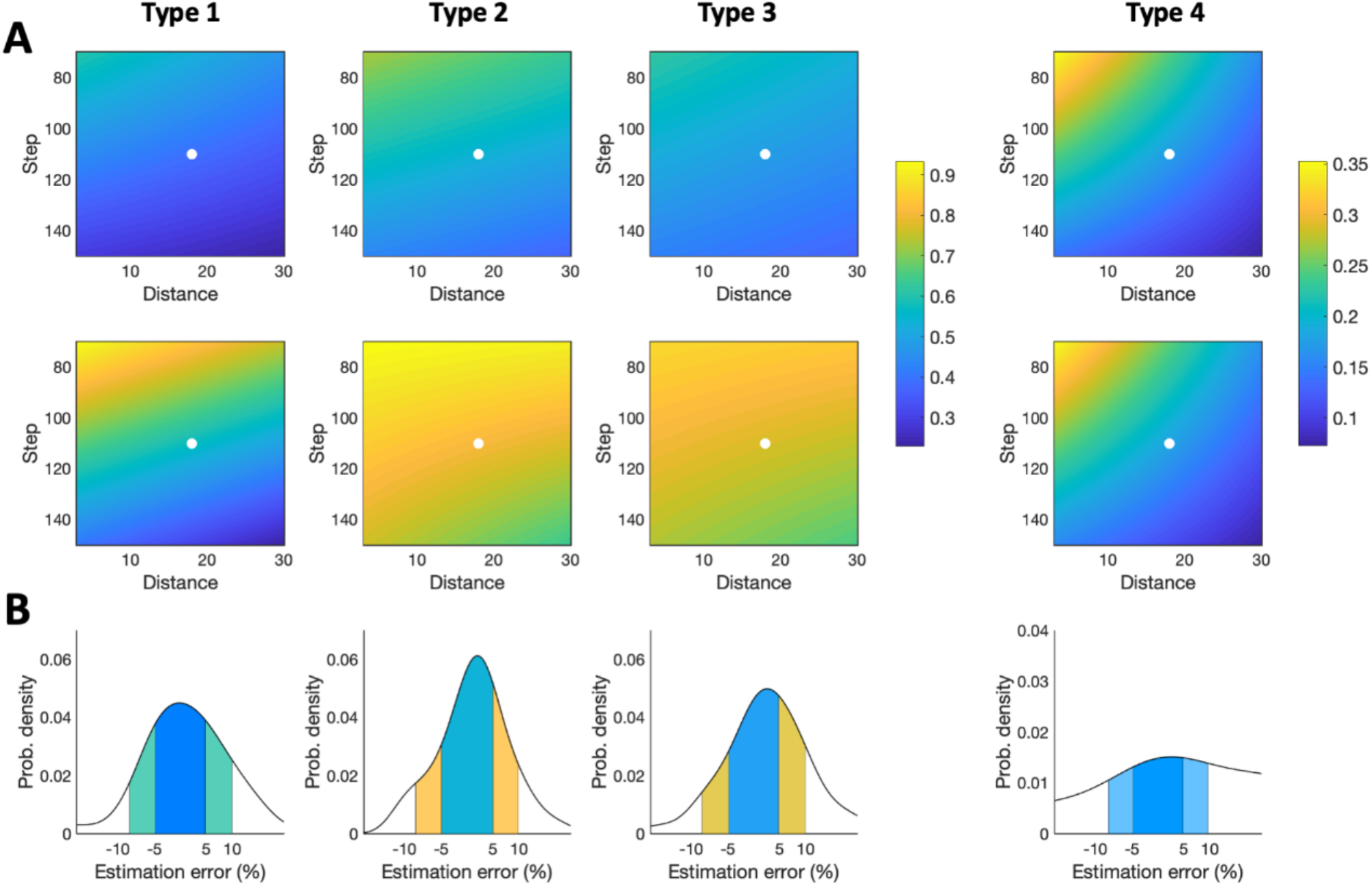
Error probability distribution for axon length estimation by planes-based stereology. **A**. Probability of ±5% (first row) and ±10% (second row) axon length estimation error for different combinations of distance and step, and for axon types 1 to 4 (columns); probability is marked by color gradient, from 0.25 to 0.92 (see color bar). Probabilities for types 1, 2, and 3 are plotted with the same colored-scale. White dots point to the cases for distance = 18 *μ* and step = 110 *μ*. Values were obtained through linear interpolation (Adjusted R-squared for 5% error: Type 1 = 0.80, Type 2 = 0.86, Type 3 = 0.81, Type 4 = 0.91; Adjusted R-squared for 10% error: Type 1 = 0.83, Type 2 = 0.83, Type 3 = 0.85, Type 4 = 0.93). **B**. Error probability distribution for the particular cases detailed in A. The areas below the curves are the probabilities shown in A and pointed with the white dots.

### Comparison of the three length estimation methods in a practical implementation in the laboratory

We measured and estimated the axonal length of three neurons corresponding to the first three axonal types considered in the study (no type 4 neurons were analyzed) by implementing in the laboratory the same approaches that were used in the computational analysis: 1) direct 3D reconstruction of the axon in Neurolucida, 2) the projection-based method performed with a camera lucida, and 3) sampling with virtual planes (See Material and Methods). Results about expected errors comparing the axonal length obtained by the three methods are summarized in **Table 1**. Although the projection-based method proved to be a robust and accurate method in the computational analysis, its practical implementation resulted in an expected error that was higher than the one obtained with stereology (11.45 vs 6.28% mean expected error). The possible sources of bias behind this difference will be discussed later.

**Table 1.**
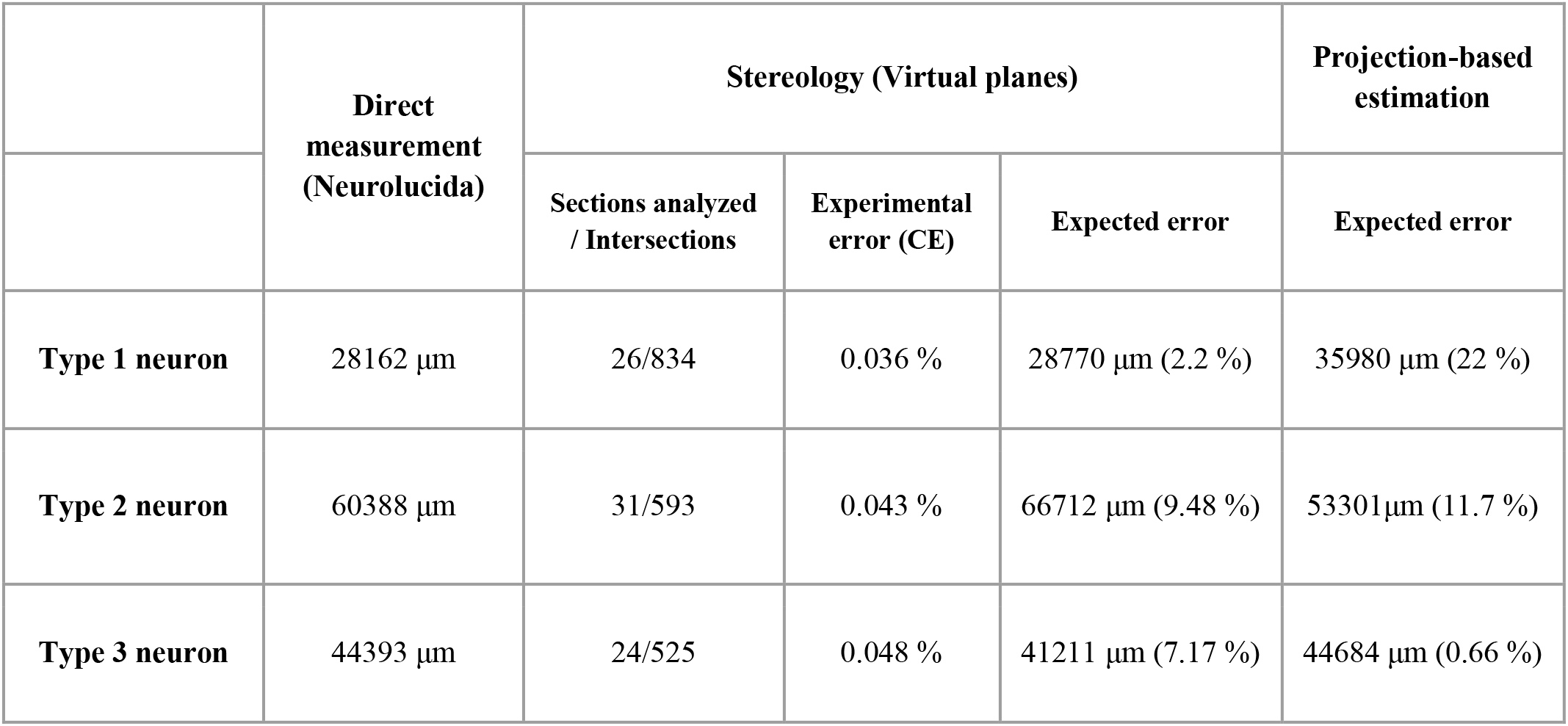
Differences between the three axonal length estimation methods after their practical implementation. Stereological parameters: step length X-Y: 75 μm, planes separation: 5 μm; CE: coefficient of error.

## DISCUSSION

We have directly tested and compared, for the first time, the accuracy and reliability of three indirect methods for length estimation of individually labeled axons visualized on serial tissue sections: 1) stereological sampling with virtual spheres [27]; 2) stereological sampling with isotropic virtual planes [26]; 3) and the correction for 2D-projected length in camera lucida drawings [22]. Using the length derived from each neuron’s 3D reconstruction as true value, we estimated each method’s accuracy or “trueness” as the closeness of agreement between the reference values and the test results. To determine if these three methods yield equally consistent results across a wide range of branching complexity and dispersion, we applied them on axons of four different cell types [36]. This analysis was performed first computationally, modeling the methods in Matlab and testing them against a dataset of mouse brain neurons from NeuroMorpho.org, and then on a number of single, thalamocortical cells labeled in serial brain tissue sections from our lab. We conclude that estimations based on stereological sampling with virtual planes or with the 2D projection-based method are a feasible and efficient alternative to complete tracing to estimate axonal length, and that they could be reliably applied to long-range projection neurons. The performance of the three methods is largely consistent over a wide range of axon branching complexity and dispersion.

The projection-based method has been used to estimate the axonal length of single thalamocortical and nigrostriatal cells labeled on histological sections [2, 7, 22]. This method consists in tracing the axons present in successive sections (using camera lucida or directly from scanned images) and then multiplying the 2D-length measurements by a mathematically-derived factor of π/2 to estimate the orthogonal length in Z-axis (**Fig. 2**). However, the actual suitability of this factor has never been tested against a large number of morphologically diverse neurons. Importantly, for this factor to be accurate, axonal length has to be isotropically distributed (with uniformly distributed orientation, [22]), something that cannot be known beforehand. Here, we used a dataset of 951 neurons of different morphologies to determine the best-fit factor, which for all neuron types (around 1.27 for types 1-3). The results of this estimation were consistent across the three orthogonal planes for every class but type 4 (local) neurons, which showed higher errors, presumably due to their smaller size and anisotropic distribution. Thus, our computational approach demonstrates the robustness of the projection-based method independently of the orientation of the plane of section, which provides accurate length estimations with a large sample of heterogeneous axonal architectures.

Design-based stereological methods can also be applied to the specific problem. However, careful selection of sampling parameters is key to make the process efficient while keeping a reasonable accuracy. As a general rule, the more exhaustive the sampling scheme (i.e., counting more intersections between the probe and the object of interest), the more precise the stereological estimation, which will in turn produce lower coefficients of error [38]. Accuracy, on the other hand, depends on the randomness of the sampling process. Although both virtual planes and spheres are based on the same principle [39], the computational results proved the first to be more accurate than the last due to restrictions imposed by the tissue thickness on the sphere’s diameter, which resulted in an inefficient and biased sampling and, consequently, a low accuracy. In this sense, large spheres (50 μm in diameter) would not get enough intersections to produce mean absolute errors below 10% when considering sections 50 μm-thick. In the case of virtual planes, accuracy depended on both the XY step (or distance between sampling boxes) and the inter-plane distance, and the results differed slightly between neuron types: because of their smaller size and their focal and dense architectures, type 1 neurons required a more intense sampling than neurons in types 2 and 3 to achieve similar mean absolute errors. Also, the range of mean absolute errors for the virtual planes method was significantly smaller than that for virtual spheres (5-12% vs 10-60%). Thus, although both virtual planes [29, 30] and virtual spheres [31–33] have been applied to estimate length in axonal arborizations arising from neuronal populations, when considering single cells in serial sections, the use of virtual planes would be preferable [34].

To corroborate the observations made with the models, the projection-based method and the stereological sampling with virtual planes were tested against three sets of tissue sections, each set containing one single thalamocortical cell of types 1-3 that had been manually reconstructed in Neurolucida. Type 4 neurons were excluded, and virtual spheres were not considered for this test because of the inaccurate results they displayed in the previous computational analysis. Virtual planes were applied using a selection of parameters derived from **Figs. 7** and **8**, and the results showed an average expected error of 6.28%, well within the range of what we had previously determined. In addition, sampling error coefficients were also small. The projection-based method, however, didn’t translate so well to the real test cases, and its practical implementation resulted in an error that was higher than expected (11.45% in average, but as high as 20% for type 1). We hypothesize that this difference might be attributable to 1) practical biases related to the process of manual tracing with the camera lucida and 2) that this randomly-picked neuron overlaps types 1 and 4. Regarding the first bias, overlapping axons could occlude each other during sampling (especially in dense and focal type 1 neurons). Thus, despite the robustness shown in the computational analysis, the accuracy of the projection-based method suffered when it was implemented onto real test cases.

Ultimately, the selection of the most suitable method may involve other practical factors, summarized in **Table 2**. In terms of economic cost, for example, the most accessible of the three methods is the projection-based approach, since it only requires a camera lucida/imaging system attachment for a microscope and a computer system with basic graphic design software. If a higher throughput and accuracy is required, the microscope could then be substituted with a slide scanner. However, this would raise the cost and make it similar to that of the specific software tools required to carry out design-based stereology (VIS or Stereo Investigator) or 3D, manual, direct reconstruction (Neurolucida). The projection-based approach requires less training, and also produces a 2D model of the neuron. 3D tracing is also relatively easy to learn and implement, whereas design-based stereology is the most complex of the three, requiring extensive know-how about sampling design or the advice of an expert. In this regard, **Figs. 6** and **7** can provide some insights that help to design the stereological procedure when using either spheres or planes probes. Thus, the choice of the adequate parameters, i.e. distance between probes and sampling intensity, will depend on the type of neuron analyzed, and aims to get the highest efficiency, which means the less analysis time to get a low error estimation (around 5%). In the case of virtual planes, type 1 neurons, which normally have short and focalized axons, need more sampling intensity than types 2 and 3 to get the same error estimation (comparison between panels 7A and 7B for the same parameter values); the opposite happens with type 4 neurons in virtual spheres, that requires less effort (less sampling intensity) than the other neuron types to get the same error estimations.

**Table 2.** Comparison of methodological approaches to measure or estimate axonal length in single neurons. The advantages and disadvantages from each of these three methods have been summarized in terms of: cost, training time, time of analysis, possible biases, the possibility of extracting additional information beyond the axonal length, and the error range computationally obtained for each neuron type and method (excluding type 4, see Figures 5–8). Different colors and +, ++ or +++ stand for different levels in each of the parameters considered. Cost: ++, affordable according to average European grant support. Training requirements: +, one day of training approximately; +++, an expert needed for sampling design/some days of training. Analysis time (per neuron): +, around 10-15 h; ++, between 50-100 h. Method biases: +, highly automated and controlled sources of bias, +++, multiple sources of biases.

| Method |  | Key system requirements | Cost | Training requirements | Time | Method Biases | Expected error range | Other Outputs |
| --- | --- | --- | --- | --- | --- | --- | --- | --- |
| **3D reconstruction**<br>(direct measurement) | **Neurolucida** | XYZ stage Neurolucida | ++ | ++ | +++ | ++ | Reference value | 3D model |
|  | **Semi-automatic reconstruction (MouseLight)** | Two-photon microscope with vibratome | +++ | ++ | + | + |  |  |
| **Projections** (model-based estimation) | **Correction of 2D length** | Camera lucida | + | + | ++ | +++ | Type 1: [1.6, 6.9]<br>Type 2: [1.0, 4.7]<br>Type 3: [1.1, 6.2] | 2D model |
|  |  | Slide-scanner | ++ | + | + | ++ |  |  |
| **Stereology** (design-based estimation) | **Space Balls** | XYZ stage Stereology software | ++ | +++ | + | + | Type 1: [10.7, 56.8]<br>Type 2: [9.7, 52.4]<br>Type 3: [15.3, 64.1] | None |
|  | **Virtual Planes** | XYZ stage Stereology software | ++ | +++ | + | + | Type 1: [4.7, 12.9]<br>Type 2: [4.0, 8.6]<br>Type 3: [5.0, 9.2] |  |

With regards to efficiency, the three methods have in common the same disadvantage: they require sparsely-labeled brains (i.e., containing only a few labeled cells that don’t overlap, to avoid errors in the reconstruction). Otherwise, axons from different neurons overlapping each other could be difficult or even impossible to distinguish from one another. With that in mind, the time required to quantify the axon length of a single neuron naturally depends on its size and spread, and more so in the case of manual 3D tracing. Whereas design-based stereology and the projection-based approach can analyze a large and complex mouse axon in about 8-16 hours, manually tracing that same cell in Neurolucida could take 10 times more. Still, the main advantage of direct reconstructions is that they also produce a 3D model of the neuron, with a collection of geometrical and topological parameters to be extracted from its structure, other than just length.

Finally, potential biases are associated with each specific method. In fact, not even direct 3D reconstruction-tracing methods, which we took as in this study, are totally free of them. The implementation of any of these methods, therefore, has to rely on clear and reproducible criteria, so as to reduce inter-individual differences between researchers when counting intersections, in the case of stereology, or tracing axons, in the case of reconstructions [40]. Also, magnification has to be relatively high (≥200X), so that individual branches are clearly recognizable as such. The greatest variation in the results is, perhaps, that introduced by the shrinkage suffered by the sections due to histological processing. Fixation procedures with paraformaldehyde and sucrose cryoprotection of the brain can introduce a shrinkage of around 15% in all dimensions, whereas the dehydration process with alcohol prior to tissue covering on the slide are known to introduce a shrinkage of up to 70% in the Z axis (therefore reducing the section thickness [41]). This shrinkage correction has to be properly applied in measurements derived from both Neurolucida and the design-based stereological approaches; otherwise, length will be underestimated. In the case of the projection-based approach, this correction is not necessary, given that it relies on 2D (XY) length, effectively ignoring the Z-dimension.

The current gold-standard in both resolution and throughput for this kind of studies are the large-scale pipelines developed by initiatives in the Allen Institute and the MouseLight projects [6, 19, 20], which are producing hundreds of neuron reconstructions registered on a template atlas in a semi-automatic manner. However, their specific requirements make them suitable to only a small number of highly-funded research centers. The methods discussed above, therefore, offer a reasonable alternative when working with sparsely labeled brains. Therefore, the majority of research groups without access to these tools can consider the methods discussed in this paper to fulfill their own research goals.

In conclusion, accurate measurement of individual axon length is crucial as this feature is an accessible estimation of the functional impact of the single neuron. Nonetheless, the technical and economic difficulties of direct axon length measurement urge for accurate, but less time and effort demanding length estimations. The three approaches presented here - projection-based length estimation, stereology, and direct measurements from reconstructions - are valid for generating accurate axonal length estimations of single cells. The stereological method is less prone to biases inherent to the manual drawing of fibers in camera lucida projection-based estimations. Among the two stereological approaches, the virtual planes method is more efficient and accurate than the one using spheres when dealing with single cell axons. Informed by the morphometric evidence presented here, the choice between these methods may thus largely depend on the human effort, training required for know-how acquisition, and economic cost. We therefore provide a useful guide for selecting the most appropriate individual axon length estimation method and the best parameters regarding the axonal morphology. Our results could be an important reference for neuroscientists interested in the wiring of neuronal circuits with powerful single-cell resolution methods, and in the building of brain models derived from this new anatomical knowledge.

## Acknowledgements

This work was supported by funding from the European Union’s Horizon 2020 Research and Innovation Programme (Grant Agreement No. 945539 - HBP SGA3), Spain’s MICINN (FLAG-ERA PCI2019-111900-2) to FC, Ministerio de Economía, Industria y Competitividad de España (BFU2017-88549) to FC and LP, and from Fundación Bancaria “la Caixa” (grant ID 100010434 - LCF/BQ/DE17/11600005) to SDH.

## Data availability

Neuron accession numbers and code used for running experiments, model fitting, plotting and statistical analysis are available on the free-access repository: https://drive.google.com/drive/folders/1qzwMSVEcKW_L6WZPLR9QSf7a3rtRMJAd?usp=sharing.

